# The IL-33-ILC2 pathway protects from amebic colitis

**DOI:** 10.1101/2021.06.14.448450

**Authors:** Md Jashim Uddin, Jhansi L. Leslie, Stacey L. Burgess, Noah Oakland, Brandon Thompson, Mayuresh Abhyankar, Alyse Frisbee, Alexandra N Donlan, Pankaj Kumar, William A Petri

## Abstract

*Entamoeba histolytica* is a pathogenic protozoan parasite that causes intestinal colitis, diarrhea, and in some cases, liver abscess. Through transcriptomics analysis, we observed that *E. histolytica* infection was associated with increased expression of IL-33 mRNA in both the human and murine colon. IL-33, the IL-1 family cytokine, is released after cell injury to alert the immune system of tissue damage during infection. Treatment with recombinant IL-33 protected mice from amebic infection and colonic tissue damage; moreover, blocking IL-33 signaling made mice more susceptible to infection and weight loss. IL-33 limited the recruitment of inflammatory immune cells and decreased the pro-inflammatory cytokine IL-6 in the colon. Type 2 immune responses, which are known to be involved in tissue repair, were upregulated by IL-33 treatment during amebic infection. Interestingly, administration of IL-33 protected RAG2^-/-^ mice but not RAG2^-/-^γc^-/-^ mice, demonstrating that IL-33 mediated protection occurred in the absence of T or B cells but required the presence of innate lymphoid cells (ILCs). IL-33 induced recruitment of ILC2 but not ILC1 and ILC3 in RAG2^-/-^ mice. Adoptive transfer of ILC2s to RAG2^-/-^γc^-/-^ mice restored IL-33 mediated protection. These data reveal that the IL-33-ILC2 pathway is an important host defense mechanism against amebic colitis.

## Introduction

Diarrhea-related illnesses are one of the major causes of death worldwide in children under five years of age, resulting in approximately 500,000 mortalities annually ^1^. *Entamoeba histolytica* is a pathogenic protozoan that is the primary causative agent of amebic dysentery. Diarrhea-associated amebic infections are correlated with growth faltering in pre-school aged children ^2^. In addition to diarrhea, infections by *E. histolytica* can also cause intestinal colitis and liver abscess, and this parasite accounts for 55,000 deaths worldwide annually ^3^. Currently, the primary treatment for amebiasis is metronidazole, a drug which, in addition to toxic side effects, has been insufficient to fully eliminate parasites from the gut ^4^. In an effort to find an effective therapeutic intervention, we sought to characterize the host immune response to amebic infection.

Host factors, along with parasite genetics and environmental factors, are likely responsible for heterogeneous outcomes of *E. histolytica* infection ^5^. Once inside the colon, amebic trophozoites can attach to intestinal epithelial cells to cause cell death and tissue damage^6^. Intestinal tissue damage by *E. histolytica* is associated with the production of pro-inflammatory cytokines, including IL-1β, IL-6, IL-8, IFN-γ, and TNFα ^7^. These pro-inflammatory cytokines may in part affect susceptibility by regulating the recruitment of neutrophils and macrophages, cells that have been shown to be critical in protection from amebiasis in a mouse model ^8^. For example, IFN-γ or TNFα treated macrophages have been shown to have potent amebicidal activity *in vitro*^9^. In children, increased production of IFN-γ by peripheral blood mononuclear cells was associated with decreased susceptibility to amebiasis ^10^. In contrast, excessive pro-inflammatory responses could be deleterious and cause host tissue damage. For example, increased TNFα in children was associated with amebic diarrhea, and blocking TNFα with neutralizing antibody was protective in a mouse model of amebiasis ^11,12^. A type 2 cytokine, IL-25, mainly released from intestinal tuft cells, was shown to protect from amebic colitis and downregulated pro-inflammatory cytokines including TNFα, IL-17a, and IL-23 ^12,13^. IL-25 treatment increased the expression of type 2 cytokines IL-4 and IL-5 and promoted the recruitment of eosinophils ^12^. Thus, complex interdependent collaborations between intestinal epithelial cells and immune cells dictate the outcome of *E. histolytica* infection. Understanding these interactions is critical to finding an effective intervention for amebiasis.

To identify host immune pathways modulated in human colonic tissue upon amebic infection, we performed an unbiased transcriptomics analysis. We observed that an IL-1 family cytokine IL-33 was upregulated in human and mouse colon. This was of interest because a number of recent studies have revealed the role of IL-33 mediated protection from experimental and infection-induced colitis in mouse models ^14–16^. Similar to IL-25, IL-33 activates type 2 innate lymphoid cells (ILC2s) and induces the expression of type 2 cytokines, and may even be a more potent inducer of type 2 immune responses ^13,14,17^. However, the role of IL-33 and ILC2s in amebic colitis has not previously been studied. Using a mouse model of amebiasis, we demonstrate here that IL-33 protected from amebic infection and colonic tissue damage. IL-33 treatment dampened overall inflammation, with decreased IL-6, CD45+ immune cells, and Ly6C^hi^ inflammatory monocytes in lamina propria. We observed that IL-33 did not require the presence of T and B cells to confer protection from amebic colitis, however did require innate lymphoid cells type 2 (ILC2s).

## Results

### Amebic infection induces IL-33 expression in humans and in the mouse model

To understand the host immune response to amebic infection, we investigated differential gene expression in human colonic biopsies via Affymetrix microarray ^18^. Samples were collected from 8 individuals at the time of infection and during convalescence (60 days after recovery). One of the significantly upregulated genes was *Il33* (Figure 1a), encoding for the cytokine IL-33, which is known as a nuclear alarmin and an inducer of type 2 immune responses. IL-25, an IL-17 family cytokine and also an inducer of type 2 immune response, was previously shown to be protective to amebic colitis ^12^. Hence, we sought to test whether IL-33 had any role in disease protection via type 2 immunity. We first tested if IL-33 was similarly increased in the colon upon *E. histolytica* infection in the mouse model of amebiasis. IL-33 mRNA and protein were measured in C57BL/6J and CBA/J mouse strains. Wildtype (WT) C57BL/6J mice are resistant to amebic colitis and start clearing the infection as early as 12 hours post-challenge ^19^. At 12 hours post-challenge, mice that remained infected had significantly higher expression of *Il33* mRNA compared to sham-challenged mice and mice that cleared the infection (Figure 1c). Unlike C57BL/6J mice, CBA/J mice are susceptible to amebic colitis and remain infected for several weeks and acquire colitis pathologically. On day 3 post-challenge, IL-33 protein was 1.6 times higher in colonic tissue of CBA/J mice compared to sham challenged mice (Figure 1d). Together, this indicated that our mouse model of infection similarly resulted in IL-33 induction as in humans.

**Figure 1:**
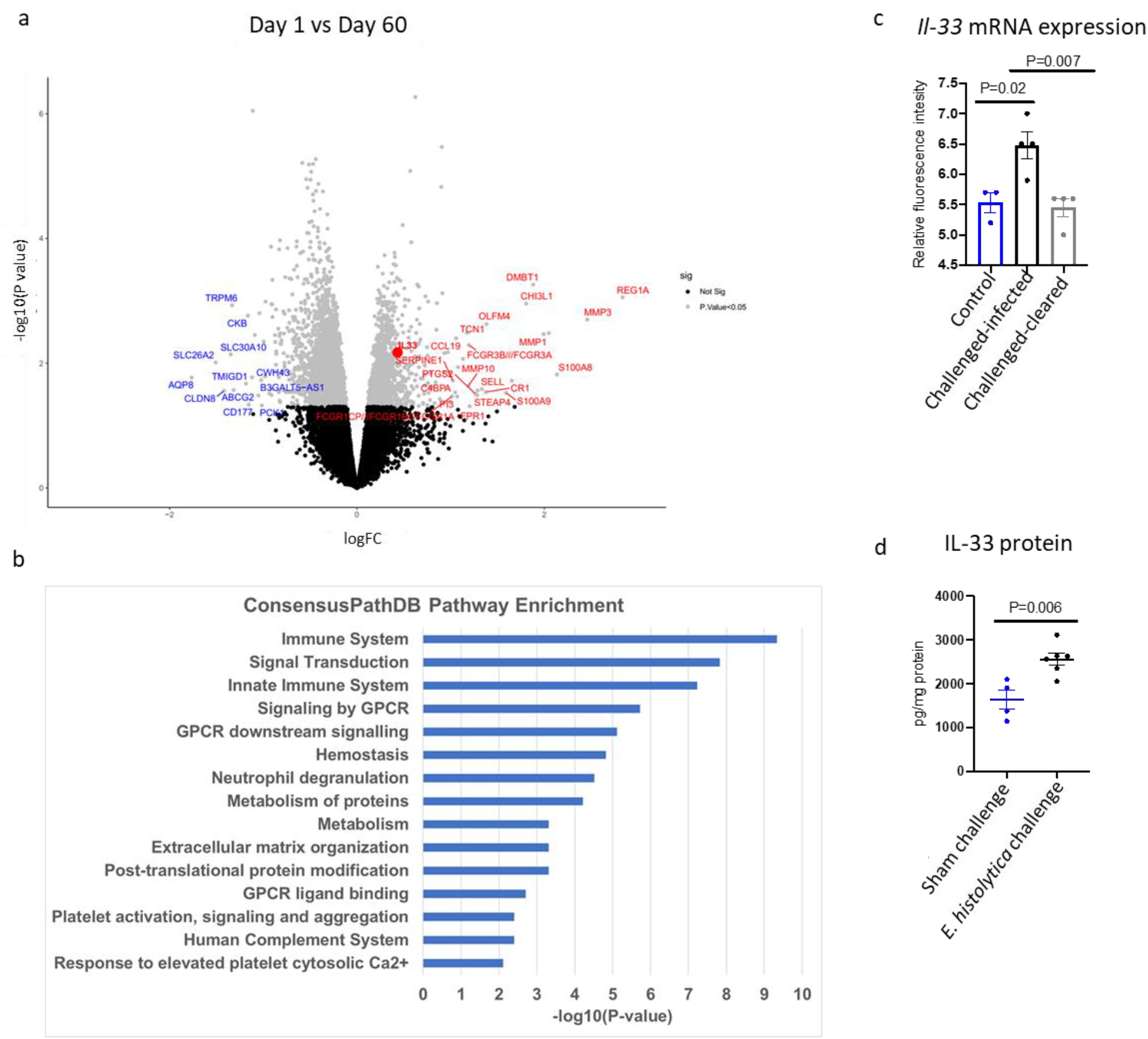
Amebic infection induces IL-33 expression in the colon of humans and mice and upregulates host immune-related pathways in colonic tissue. **a-b** Transcriptomic analyses were performed on colon biopsies collected at acute amebic infection (at day 1) and convalescence (at day 60) of 8 patients. **a** Volcano plot shows IL-33 (red dot) among upregulated genes. **b** Pathway enrichment analysis of upregulated transcripts (logFC>=0.4) using Consensus PathDB tool. **c-d** IL-33 mRNA and protein were measured in mouse cecal tissue. **c** *Il-33* mRNA expression by microarray from C57BL/6J mice, n = 3-4 per group. **d** IL-33 protein by ELISA in cecal tissue lysate from CBA/J mice, data representative of two independent experiments, n =4-5 per group. Statistical significance was determined by one-way ANOVA and unpaired t test. Error bars indicate SEM.

The upregulated genes (logFC>0.4) from the human colon transcriptome were analyzed with the Consensus PathDB pathway enrichment tool. Innate/adaptive immunity and G-protein coupled receptor signaling were among the upregulated pathways (Figure 1b). Interestingly, pathways involved in tissue repair (hemostasis, extracellular matrix organization) were also significantly upregulated. These results suggested that increased IL-33 might be involved in the induction of barrier repair mechanisms to protect gut tissue from amebic colitis.

### IL-33 protects from amebic colitis

As infection increased IL-33 in both humans and in mice, we utilized the mouse model of amebic colitis to investigate if IL-33 protected from amebiasis. CBA/J mice were treated with 0.75 µg of recombinant IL-33 intraperitoneally for eight days (from day-3 to day+4 of the amebic challenge). On day+5 of the challenge, mice were sacrificed to collect cecal content and cecal tissue to determine the infection rate and epithelial damage. Treatment with IL-33 significantly protected mice from infection and weight loss (Figure 2a, 2b, and 2c). Hematoxylin and eosin (H&E) staining showed that the IL-33-treated group had a more intact epithelium compared to the PBS treated group (Figure 2d). These data supported a role of IL-33 to clear ameba and protect and/or repair the colonic epithelial layer from damage.

**Figure 2:**
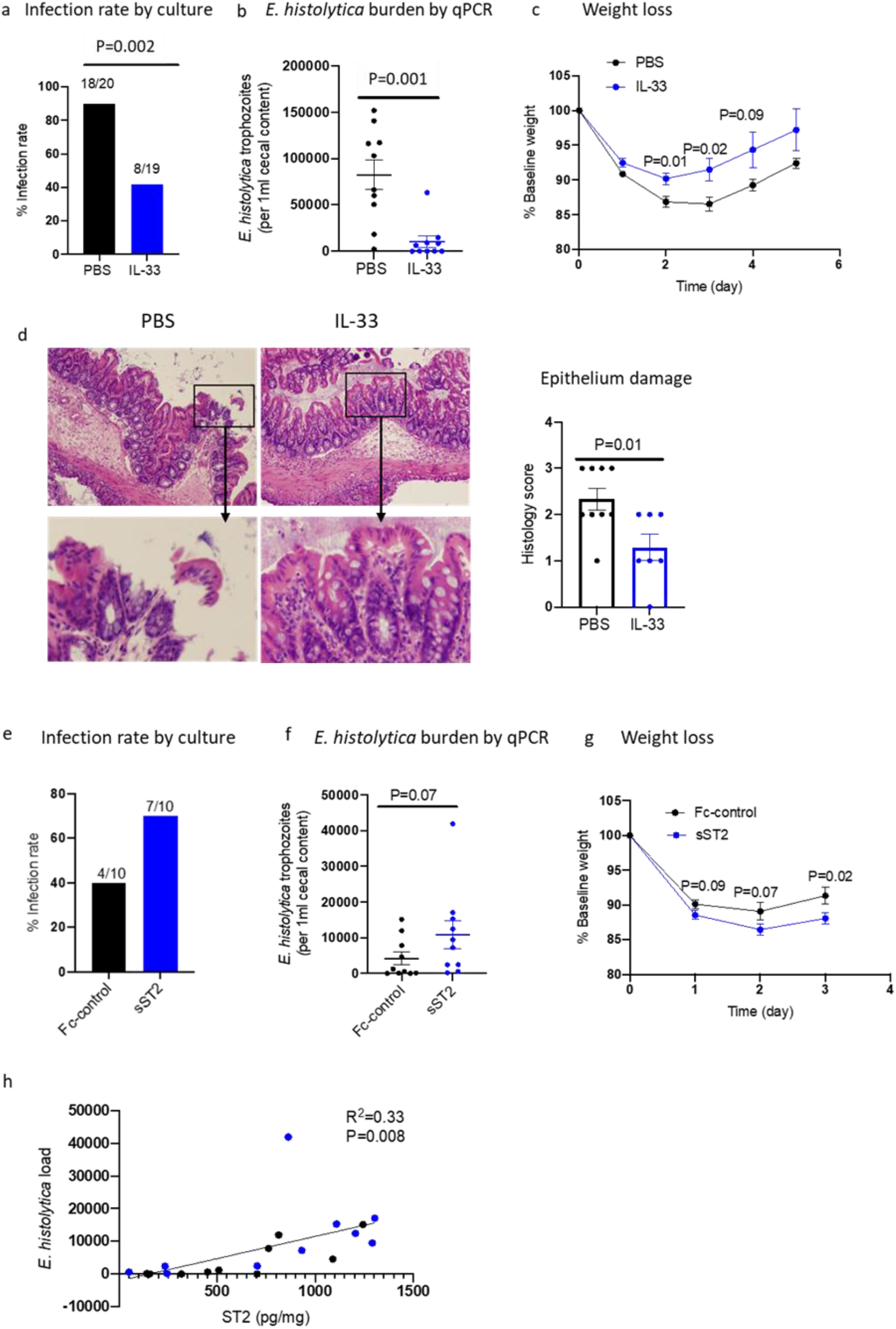
IL-33 protects from amebic infection and epithelial tissue damage in mice. **a-d** CBA/J mice were injected intraperitoneally with 0.75 µg of IL-33 or PBS each day for 8 days. On day 4 mice were challenged with *E. histolytica* trophozoites. Cecal content and cecal tissue were harvested on day 9 (day 5 post *E. histolytica* challenge). **a** Infection rate measured by amebic culture from cecal content. **b** *E. histolytica* DNA, measured by qPCR. **c** Weight loss. **d** Epithelial damage scored by H&E staining of cecal tissue. **e-h** Impact of soluble ST2: C57BL/6J mice were intraperitoneally injected soluble ST2 (IL-33 receptor). **e** infection rate by amebic culture from cecal content. **f** *E. histolytica* DNA by qPCR. **g** Weight loss. **h** Correlation of amebic DNA load and ST2 concentration in cecal tissue (blue dots represent mice in soluble ST2 group and black dots represent control group). **a** Data pooled from 2 independent experiments (n=9-10). **b-d** Data representative of 2 independent experiments (n=9-10). **e-h** Data pooled from two independent experiments (n=5-10). Statistical significance was determined by Fisher’s exact test and unpaired t test. Error bars indicate SEM.

We endeavored to know whether endogenous IL-33 signaling had a role in this protection. IL-33 signals through binding the IL-1 receptor-like protein ST2 ^20^. The IL-33 receptor ST2 can be found in both a cell surface-bound form as well as a soluble form. The soluble form of ST2 can bind with IL-33 and act as a negative regulator of the IL-33 signaling pathway ^21^. Downregulation of IL-33 signaling by adding soluble ST2 made mice more susceptible to infection (Figure 2e and 2f). Soluble ST2 treated mice lost significantly more weight than vehicle-treated mice upon challenge with amebic trophozoites (Figure 2g). Notably, the concentration of ST2 in colonic tissue was positively correlated with the colonic burden of *E. histolytica* (Figure 2h). Together, these data demonstrated that endogenous IL-33 protected from colonic amebiasis.

### IL-33 regulates the type 2 immune response and dampens colonic inflammation upon amebic infection

IL-33 has been shown to induce a type 2 immune response in intestinal tissue during helminth and bacterial infections ^15,22^. We hypothesized that IL-33 may be protecting in our model by promoting a type 2 immune response. To examine this, we used RT qPCR to examine if the transcription of cytokines that are associated with a type 2 response, *Il-4, Il-5*, and *Il-13*, were altered. While IL-33 treatment increased the transcription of IL-5 and IL-13, we did not observe any significant change of IL-4 transcript levels (Figure 3a). Others have found that IL-33 can induce IL-13 secretion from innate lymphoid cells leading to differentiation of goblet cells ^24^. Intestinal goblet cells secret a mucus-gel that is known to contribute to the clearance of amebic trophozoites ^25,26^. Thus, we sought to determine if IL-33 was altering goblet cells in our model. Examining PAS-stained fixed tissues, we found that goblet cells were increased in the IL-33 administered group (Figure 3b). The primary structural component of the mucus-gel is the MUC2 mucin. In concordance with our histological analysis, we also found that IL-33 treated mice had higher expression of *Muc2* mRNA compared to PBS treated mice (Figure 3c).

**Figure 3:**
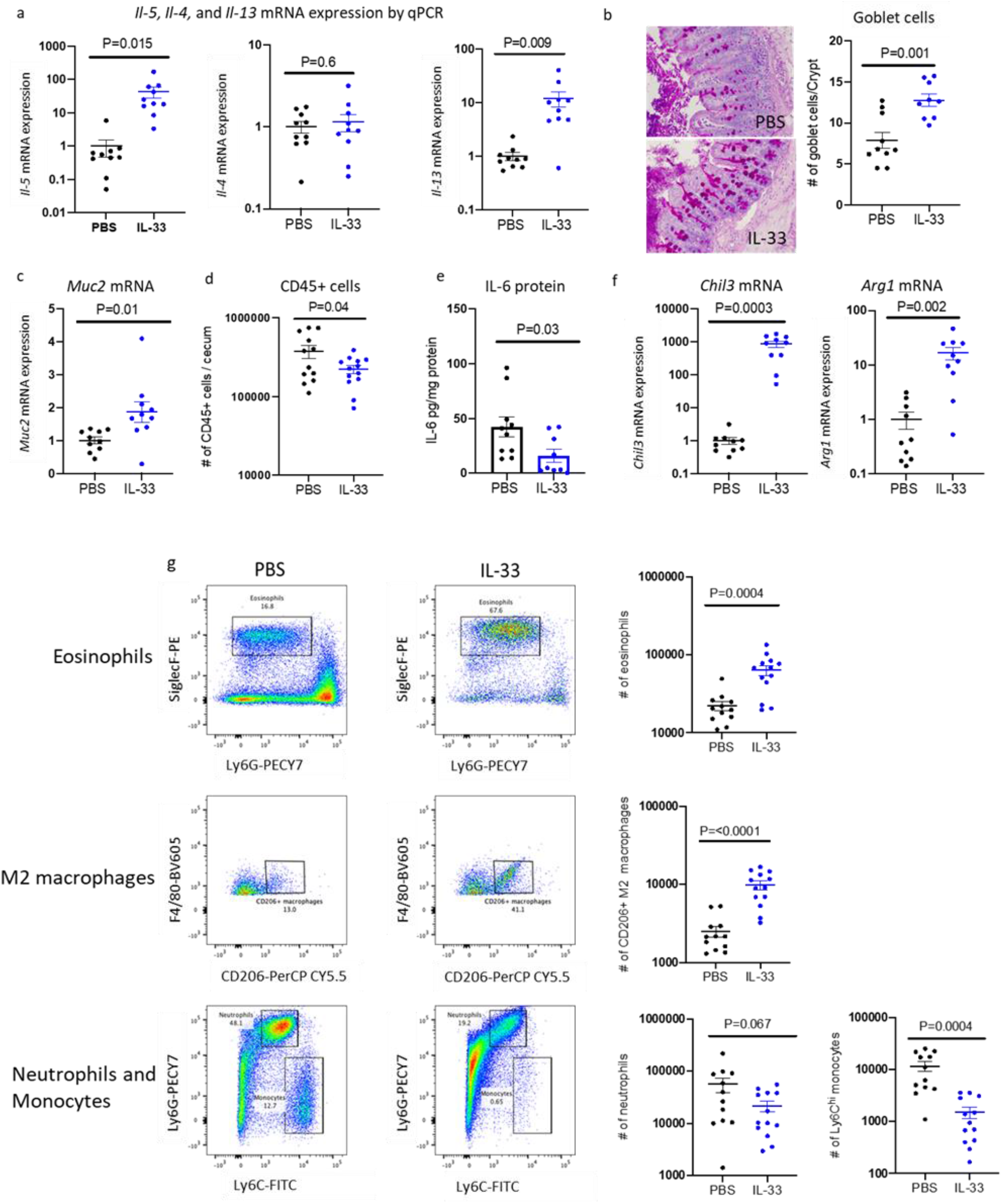
IL-33 upregulates type 2 immune response and dampens inflammation during amebic colitis. **a-g** Cecal tissue was collected on day 5 from IL-33 or PBS treated mice. **a** mRNA expression of type 2 cytokines. **b** PAS-stained goblet cells by microscopy. **c** *Muc2* mRNA expression. **d** Total CD45+ cells in cecal tissue by flow-cytometry. **e** IL-6 protein by ELISA from cecal lysate. **f** *Chil3* and *Arg1* (M2 macrophages marker) mRNA expression. **g** Flow cytometry gating and immune profiling of myeloid cells. **a-f** Data representative of 2 independent experiments (n=5-10). **g** Data pooled from 2 independent experiments (n=5-10). Statistical significance was determined by unpaired t test. Error bars indicate SEM.

IL-5 is crucial in the maturation and release of eosinophils ^23^. Coordinately, flow-cytometric analysis of cecal tissue revealed that administration of IL-33 resulted in eosinophilia during amebic infections (Figure 3g, top panel). Since IL-33 treatment upregulated colonic eosinophils, we asked if IL-33 mediated protection was dependent on eosinophilia. To test that, we depleted eosinophils in IL-33 treated mice with anti-SiglecF monoclonal antibody (Supp. Figure 1a). Treatment with anti-SiglecF successfully depleted eosinophils but not alternatively-activated macrophages or monocytes in colonic tissue (Supp figure 1e, 1f, and 1g). There was not an increase in infection rate with anti-SiglecF administration, and amebic load and weight loss were not different between the treatment and control groups (Supp. Figure 1b, 1c, and, 1d). These data suggested that IL-33 mediated protection from amebic colitis was not conferred by eosinophils.

In addition to regulating the type 2 response, IL-33 has been shown to dampen the inflammatory response during intestinal infection ^15^. In our model, there was a significantly decreased number of CD45+ cells in cecal tissue in IL-33 treated mice (Figure 3d). In addition to total CD45+ cells, the IL-33 treated group had a significantly decreased number of Ly6C^hi^ inflammatory monocytes (Figure 3g, lower panel) as well as the pro-inflammatory cytokine IL-6 (Figure 3e). There was a trend of lowered Ly6G+ neutrophils, although that was not statistically significant (P=0.067) (Figure 3g, lower panel). Interestingly, IL-33 treatment expanded the number of alternatively-activated macrophages, which are thought to be anti-inflammatory (Figure 3g, middle panel). Markers of anti-inflammatory macrophages such as Chil3 and Arg1 were also found to be significantly higher in IL-33 treated mice (Figure 3f). Together these data demonstrated that IL-33 induced a type 2 immune response and diminished inflammations during amebic colitis.

### IL-33 mediated protection from amebic colitis requires ILCs but not T and B cells

IL-33 has been shown to promote the function of regulatory T cells as well as ILC2s in intestinal tissue to protect from barrier disruption ^16,27^. Moreover, IL-33 has been shown to act on B cells to promote the production of IgA, which, in turn, protects mice from colitis and colitis-associated cancer ^28^. Hence, it was important to determine which cell types are responsible for IL-33 mediated protection from amebiasis. We treated RAG2^-/-^ mice (deficient of T and B cells) with recombinant IL-33 and observed that even in the absence of T and B cells, IL-33 still protected from amebic infection and weight loss (Figure 4a, 4b, and 4c). These data indicated that the presence of T and B cells was not necessary for this IL-33 mediated defense.

**Figure 4:**
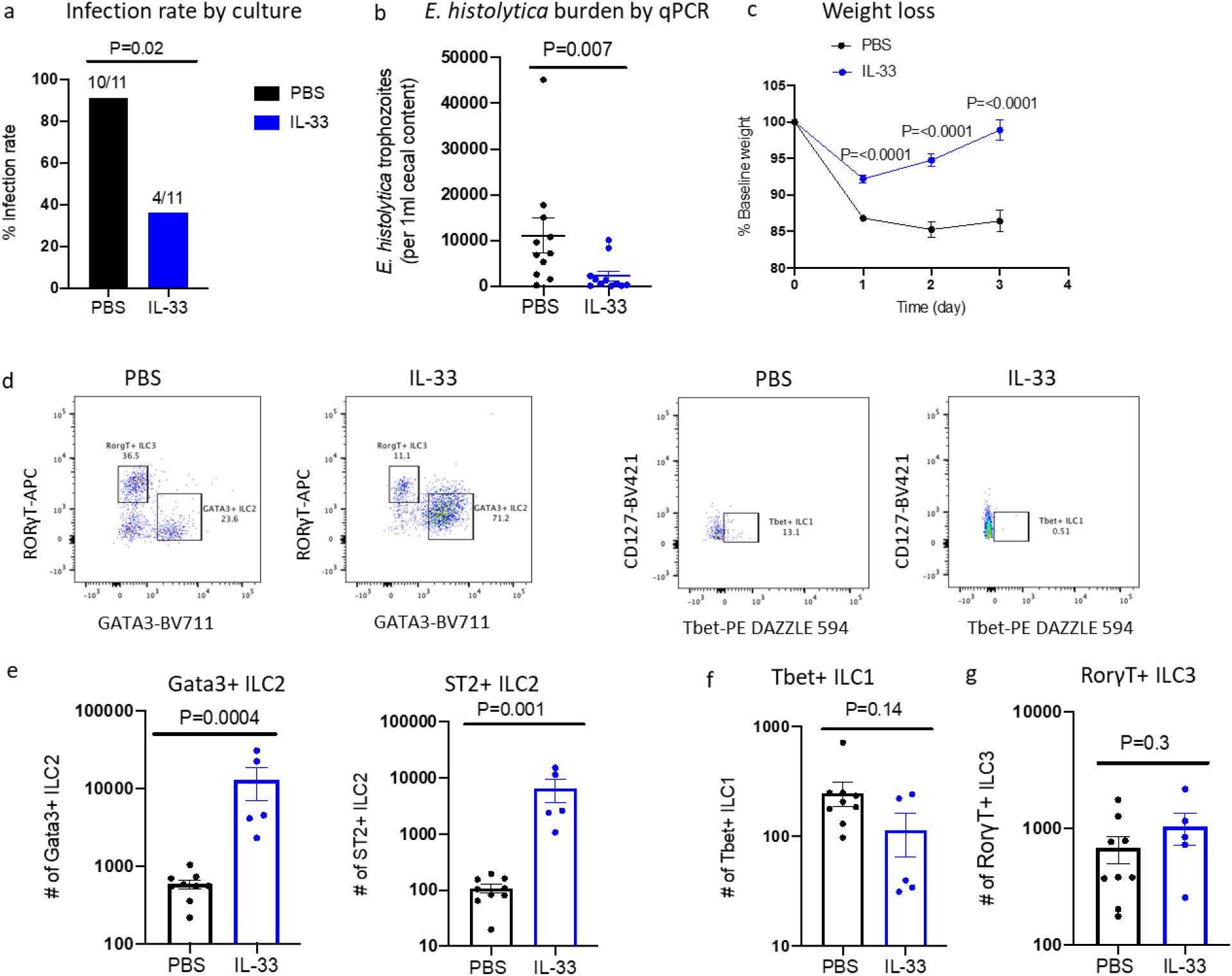
IL-33 protects RAG2^-/-^ mice and upregulates Gata3+ and ST2+ ILC2s. **a-g** RAG2^-/-^ mice (C57BL/6 background) were treated with IL-33 and PBS followed by an amebic challenge. On day 3 post-amebic challenge mice were sacrificed to collect cecal content and cecal tissue. **a** Infection rate determined by amebic culture of cecal content. **b** *E. histolytica* DNA measured by qPCR from cecal content. **c** Weight loss. **d** Flow cytometry gating to determine the total number of ILC1, ILC2, and ILC3. **e** Total number of Gata3+ and ST2+ ILC2s in cecal tissue. **f** Number of ILC1. **g** Number of ILC3. **a-c** Data pooled from two independent experiments (n=5-6). **d-g** Data representative of two independent experiments (n=5-9). Statistical significance was determined by Fisher’s exact test and unpaired t test. Error bars indicate SEM.

Interestingly, RAG2^-/-^ mice that received IL-33 had significantly more Gata3+ ILC2s in their colon compared to vehicle-treated mice (Figure 4d and 4e), while the number of ILC1s and ILC3s was not significantly different (Figure 4f and 4g). Most of the Gata3+ ILC2 also expressed the IL-33 receptor (ST2) (Figure 4e), suggesting that ILC2s acted downstream of IL-33 signaling. These data suggested a potential role of ILC2s in IL-33 promoted defense from amebic colitis.

To further delineate the importance of ILCs, we utilized a RAG2^-/-^γc^-/-^strain of mouse that does not have ILCs in addition to lacking T and B cells. While IL-33 was able to protect RAG2^-/-^mice, protection was lost in RAG2^-/-^γc^-/-^mice (Figure 5a, 5b, and 5c). In addition, upregulation of type 2 immune responses was abrogated in RAG2^-/-^γc^-/-^mice (Figure 5d and 5e). This included MUC2 mucin that was upregulated in RAG2^-/-^ mice but not in RAG2^-/-^γc^-/-^ mice upon IL-33 treatment (Figure 5f).

**Figure 5:**
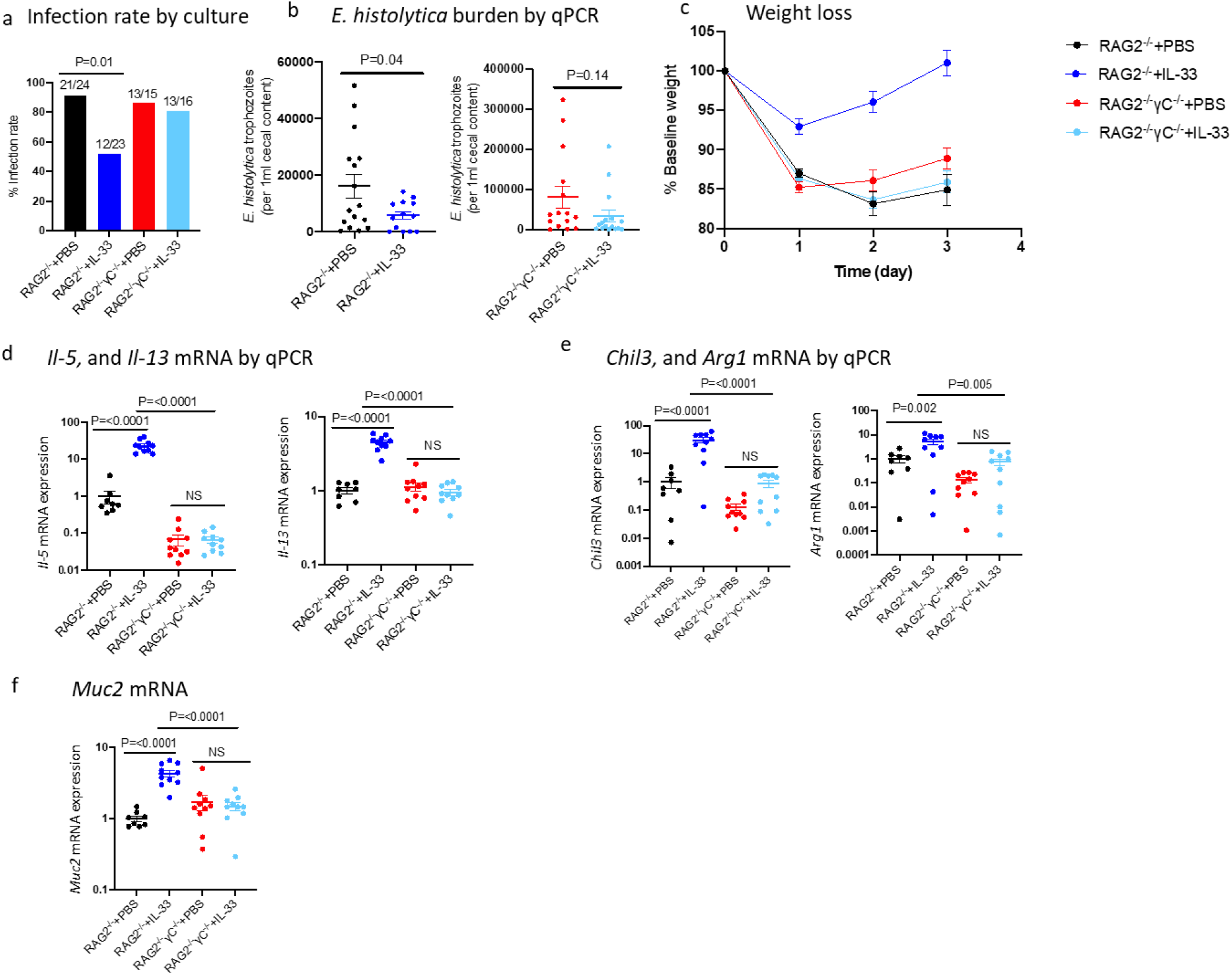
IL-33 mediated protection from amebiasis and upregulation of type 2 immune responses are lost in RAG2^-/-^γc^-/-^mice. RAG2^-/-^and RAG2^-/-^γc^-/-^ mice were treated with IL-33 or PBS and challenged with amebic trophozoites. **a** Infection rate by amebic culture. **b** *E. histolytica* DNA. c Weight loss. **d** *Il5*, and *Il13* mRNA expression. **e** *Chil3*, and A*rg1* mRNA. **f** *Muc2* mRNA. **a** Data pooled from 3 independent experiments (n=5-10). **b** Data pooled from 2 independent experiments (n=5-10). **c-f** Data representative of 2 independent experiments (n=5-10). Statistical significance was determined by Fisher’s exact test, one-way ANOVA, and unpaired t test. Error bars indicate SEM.

### Adoptive transfer of ILC2s restored IL-33 mediated protection in RAG2^-/-^γc^-/-^ mice

To determine if ILC2s acted downstream of IL-33 signaling during protection from amebic colitis, we adoptively transferred ILC2s into RAG2^-/-^γc^-/-^ mice. ILC2s were isolated from mouse spleen, mesenteric lymph node, and colonic tissue and enriched using magnetic cell separation. Isolated cells were then expanded *in vitro* and activated in the presence of IL-33 [5,13,14]. ST2+ ILC2s were flow-sorted and adoptively transferred into RAG2^-/-^γc^-/-^ mice (5×10^5^ cells/mouse) through intraperitoneal injection. Both mice receiving ILC2s and control mice were treated with IL-33 (Figure 6a). While 100 percent of the control RAG2^-/-^γc^-/-^ mice remained positive for ameba on day 3 post amebic challenge, 55 percent of RAG2^-/-^γc^-/-^ mice that received ILC2s were culture-negative (Figure 6b). *E. histolytica* DNA load was also significantly lower in ILC2s administered mice (Figure 6c). ILC2 treatment also protected RAG2^-/-^γc^-/-^ mice from weight loss (Figure 6d). The successful transfer of ILC2s in RAG2^-/-^γc^-/-^ mice was confirmed by flow cytometry analysis (Figure 6e). These data demonstrate that ILC2s acted downstream of IL-33 during protection from amebic colitis in the mouse model.

**Figure 6:**
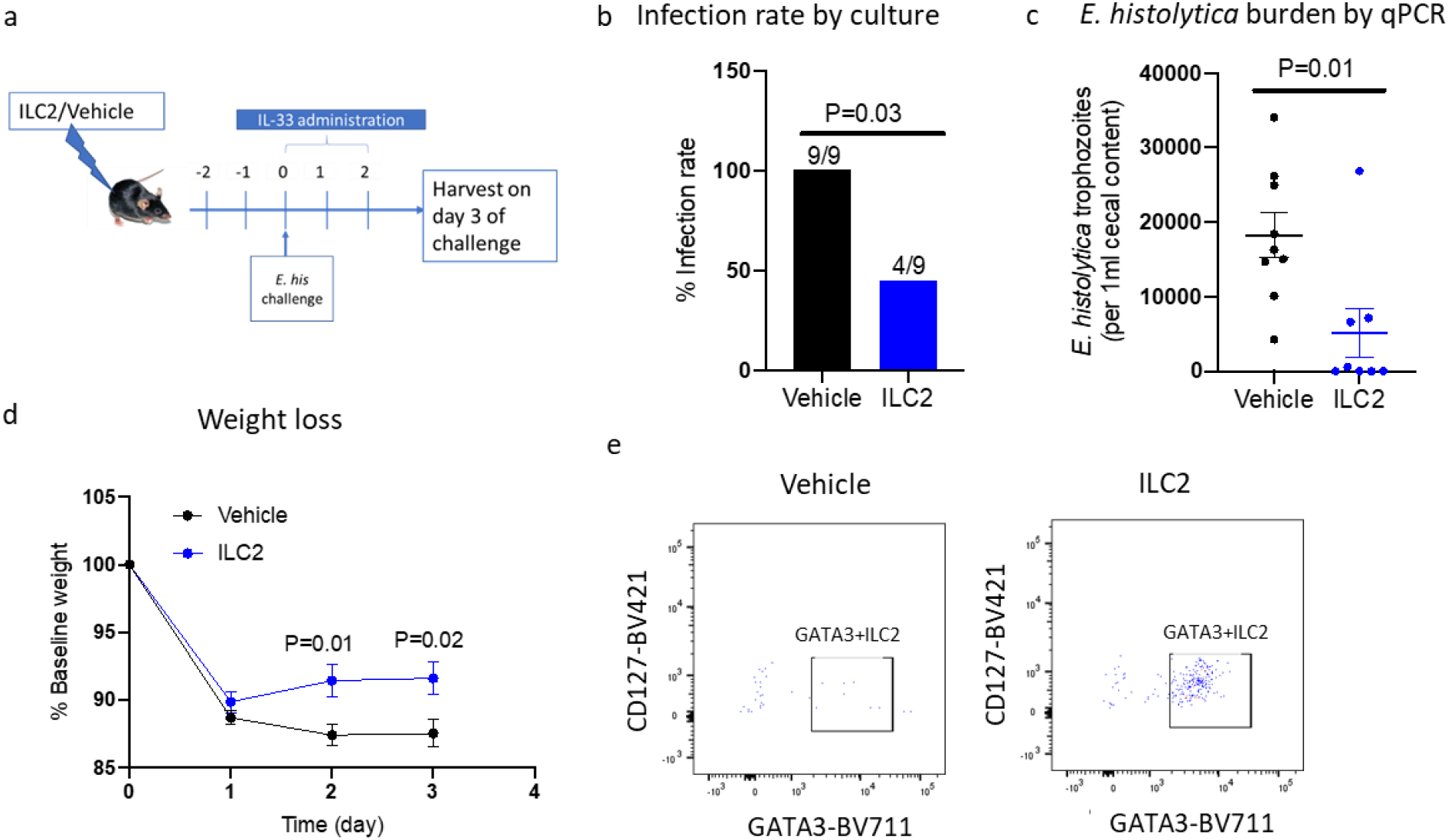
IL-33 protects ILC2 transplanted RAG2^-/-^γc^-/-^ mice from amebic colitis. **a-e** RAG2^-/-^γc^-/-^ mice were transplanted with flow-sorted ILC2s or vehicle followed by amebic challenge. On day 3 post-challenge mice were harvested and cecal tissue and cecal contents were collected. **a** Experimental outline. **b** Infection rate by amebic culture of cecal content. **c** Amebic load by qPCR. **d** Weight loss. **e** Representative flow-cytometry gating of ILC2s. **b-d** Data pooled from 2 independent experiments (n=3-6). Statistical significance was determined by Fisher’s exact test and unpaired t test. Error bars indicate SEM.

## Discussion

The most important discovery of this work is that the nuclear alarmin IL-33 protects from intestinal colitis caused by the protozoan parasite *E. histolytica*. IL-33 was found to mediate innate immunity via an ILC2-driven type 2 immune response. The upregulation of IL-33 transcripts in the human colon during amebic infection was the first indication of a potential role of IL-33 signaling in amebiasis. In the mouse model, treating with recombinant IL-33 cleared trophozoites from the mouse colon and helped in early recovery from colitis, as evidence by decreased weight loss and epithelial damage. Conversely, inhibition of IL-33 signaling by administration of its soluble receptor ST2 increased susceptibility. Finally, ILC2s were shown to be required by restoration of IL-33 mediated protection by adoptive transfer into RAG2^-/-^ γc^-/-^ mice.

IL-33 expression was previously found to be induced in inflamed colonic tissues from ulcerative colitis patients ^29^. In mouse models, colonic infections by helminths and bacteria increased IL-33 expression ^15,30^. Similarly, we observed induced IL-33 expression both in the colon of humans and mice during amebic infection. The mechanisms through which *E. histolytica* infection increases IL-33 expression are not known. Studies show that activation of TLR2/TLR6 signaling with agonist upregulates IL-33 transcripts in an NLRP3 dependent manner ^31^. Amebic lipopeptidophosphoglycan is recognized by TLR2 and TLR4 on innate and adaptive immune cells resulting in production of effector cytokines ^32^. Thus, amebic PAMPs could signal through TLR2/TLR4 to activate NLRP3-inflammasome and induce IL-33 expression. In addition, cell death is a known mechanism of IL-33 induction, and it is possible that IL-33 is upregulated by the novel means by which *E. histolytica* kills human cells called “trogocytosis” (where amebic trophozoites nibble a part of target cells resulting in cell killing and tissue invasion) ^6^.

The upregulation of IL-33 with an amebic infection that is part of the host protective response may act in part by inducing tissue repair mechanisms that enhance blood coagulation and angiogenesis in colonic tissue ^16,33^. Pathway analysis of transcript data derived from colonic biopsies of *E. histolytica* infected patients revealed that extracellular matrix organization and hemostasis were two of the top ten upregulated pathways. These pathways are involved in the initiation of tissue repair and recovery from epithelial damage. A recent study showed that IL-33 administration protected mice from colonic tissue damage and increased barrier integrity during *C. difficile* infection ^15^. In this study, we found that treatment with IL-33 reduced epithelial damage during amebiasis. IL-33 also reduced parasitic loads from the colon, likely due in part to upregulation of genes involved in innate immune signaling that were overrepresented during acute amebic infection.

IL-33 may act to protect from amebiasis in part by dampening inflammation. Inflammation and immune cell recruitment are host-protective responses towards injury and infection; however, chronic and uncontrolled inflammation could cause collateral damage to the host. With IL-33 treatment, we observed a reduced number of CD45+ cells and Ly6C^hi^ inflammatory monocytes and lower IL-6. IL-6 can induce chronic inflammatory and autoimmune responses ^34^. In addition, IL-33 skewed the immune response towards type 2 immunity, which was previously shown to be protective during helminth and bacterial infection ^15,35^. Our lab has previously shown that IL-25, an IL-17 family cytokine that can also lead to a type 2 immune response, is protective to intestinal amebiasis. Like IL-33, IL-25 defends from amebic colonization and intestinal tissue damage. IL-25 protected through induction of eosinophilia as observed depletion of eosinophils by neutralizing antibody abrogated protection. With IL-33 treatment, we also observed an increase in colonic eosinophils. Surprisingly, depletion of eosinophils did not abrogate the IL-33 mediated protection (Supplementary figure 1). These data suggest that IL-25 and IL-33 use different downstream pathways to protect from amebic colitis.

IL-33 can activate different ST2+ lymphoid and myeloid cells, including mast cells, basophils, Th2 cells, Treg cells, and ILC2s ^30^. We sought to determine if IL-33 was acting on Th2 cells or/and ILC2 cells to protect from amebic colitis by inducing a type 2 immune response. We observed that IL-33 treatment protected RAG2^-/-^ mice on a C57BL/6 background in a manner similar to WT CBA/J mice. However, this protection was lost in RAG2^-/-^γC^-/-^ (C57BL/6 background) mice. IL-33 mediated upregulation of IL-5, IL-13, Muc-2, Chil3, and Arg-1 transcripts was abrogated in RAG2^-/-^γC^-/-^ mice, but protection was restored by adoptive transfer of ILC2. These data indicated that the presence of ILC2s but not Th2 cells, was essential for the upregulation of type 2 immune responses and protection from amebic colitis.

ILCs are the innate counterpart of CD4+ T helper cells, as evidenced by their effector cytokine production and master transcription factors. Unlike T cells, they do not have antigen receptors; however, they possess receptors for cytokines and are regulated by IL-33, IL-25, and TSLP upon injury or infection ^36^. ILCs have been found to shape intestinal health in diseased or homeostatic conditions. However, the role of ILCs in amebic colitis had not been studied. We observed that IL-33 treatment increased the number of Gata3+ and ST2+ ILC2s in mouse cecal tissue upon amebic infection. Adoptive transfer of ILC2s was sufficient to restore the IL-33 mediated protection from amebic colitis in RAG2^-/-^γC^-/-^ mice. IL-33-ILC2 induced goblet cell hyperplasia has been implicated in helminth expulsion from the intestine ^37^. Additionally, IL-33 activated CD206+ type 2 macrophages were associated with tissue repair in experimental colitis ^14^. There was a significantly higher number of goblet cells and expression of Muc2 mucin in cecal tissue upon IL-33 treatment. We also observed an increased number of CD206+ alternatively-activated macrophages. Future studies should focus on delineating the contribution of IL-13 mediated goblet cells hyperplasia, IL-5, and alternatively activated macrophages in clearance of amebic infection and recovery from colonic tissue damage.

A recent study showed that ILC2s exacerbated amebic liver abscess in a mouse model through IL-5 facilitated accumulation of eosinophils in the abscesses ^38^. While type 2 immune-mediated wound healing is key to recovery from colonic tissue damage by infections, this could be deleterious in some organs, including lung and liver ^14,15,38–41^. Tissue-specific heterogeneity in ILC2s has been demonstrated, and could contribute to the apparently different roles of ILC2 in response to colitis and liver abscess due to *E. histolytica* ^42^. Single-cell RNA sequencing of ILC2s collected from amebic infected colonic tissue could further advance the understanding of downstream effector pathways involved in ILC2 mediated protection from amebic colitis and how these differ from liver abscess.

Altogether, we showed for the first time that *E. histolytica* infection upregulates IL-33 transcripts in human and mouse colonic tissue. Endogenous IL-33 signaling and exogenous treatment with recombinant IL-33 protected from amebic colitis. This IL-33 mediated defense from amebiasis was associated with the induction of type 2 immune responses and while IL-33 did not require T and B cells, the presence of ILC2s was sufficient to confer protection. This work showed that the IL-33-ILC2 pathway is important to clear parasitic loads and recover from tissue damage during colitis and could be targeted to design effective interventions.

## Methods

### Mice

Experiments were performed using sex-matched 7-10 weeks old C57BL/6J, CBA/J, Rag2^-/-^, or Rag2^-/-^γC^-/-^ mice. C57BL/6J and CBA/J mice were purchased from Jackson Labs and housed in specific pathogen-free conditions in the vivarium of the University of Virginia. Rag2^-/-^ and Rag2^-/-^γC^-/-^ mice with an excluded flora were purchased from Taconic Biosciences. Rag2^-/-^ and Rag2^-/-^γC^-/-^ mice had bedding swapped twice a week for 3 weeks for the purpose of equilibrating microbiota before experiments were performed. All the experiments were approved by the University of Virginia Institutional Animal Care and Use Committee.

### Recombinant IL-33 treatment

Mice were administered 0.75 µg of recombinant IL-33 (BioLegend, San Diego, CA) or PBS via intraperitoneal injection daily beginning at 3 days before amebic challenge up until 1 day before harvest.

### *E. histolytica* infection

Mice were challenged with 2×10^6^ *E. histolytica* trophozoites (laboratory strain HM1: IMSS) through intra-cecal laparotomy ^19^. At harvest, cecal contents were collected and cultured in complete TYI-S-33 medium ^43^. After 24 hours of culture, the presence of *E. histolytica* trophozoites in culture tubes was determined by microscopic examination to determine infectivity. Cecal contents (200 µl) were also used to extract DNA using QIAamp Fast DNA Stool Mini Kit (Qiagen, Hilden, Germany). *E. histolytica* DNA was quantified using real-time PCR (qPCR) from extracted DNA. The primers and probe used for qPCR are as follows-Eh-probe: Fam/TCATTGAATGAATTGGCCATTT/BHQ; Eh-forward: ATTGTCGTGGCATCCTAACTCA; Eh-reverse: GCGGACGGCTCATTATAACA.

### H&E staining and PAS staining

At harvest, mouse cecal tissue sections were collected and fixed into Bouin’s solution for 24 hours and then transferred into 70% ethanol. Tissue sections were then embedded into paraffin, sectioned and stained by H&E stain or Periodic acid-Schiff (PAS). Two independent blinded scorers scored the epithelium disruption (ranged from 0-3) and counted goblet cells ^44^. Goblet cell numbers were normalized to the number of crypts.

### Preparation of tissue lysate and tissue protein measurement

Dissected cecal tissue was mixed with 300 µl of lysis buffer 1 (1× HALT protease inhibitor, 5 mM HEPES) followed by bead beating for 1 minute. After bead beating, 300 µl of lysis buffer 2 (1× HALT protease inhibitor, 5 mM HEPES, 2% Triton X-100) was added with the solution and incubated on ice for 30 minutes. Solutions were centrifuged for 10 minutes at 10,000 x *g*, and supernatants were collected. IL-33 and ST2 protein concentration was measured using R&D ELISA kits (Minneapolis, MN). Protein concentrations were normalized to total tissue protein.

### RNA extraction and qPCR

RNA was extracted from cecal tissue using a RNeasy isolation kit (Qiagen, Hilden, Germany). Genomic DNA was removed using a Turbo DNA-free kit (Invitrogen, Carlsbad, CA). RNA was then transcribed to cDNA using a High-Capacity cDNA Reverse Transcription kit (Applied Biosystems, Foster city, CA). Expression of *Il-5, Il-13, Il-4, Muc2, Chil3*, and *Arg1* genes was quantified by qPCR using Sensifast SYBR & Fluorescein kit (Bioline, London, UK). Gene expression was normalized to Gapdh expression. All procedures were performed following protocols provided in the kits. Primers used for qPCR are as follows: *Il-13*: F-5′-CAGCATGGTATGGAGTGTGGACCT-3’, R-5′-ACAGCTGAGATGCCCAGGGAT-3’; AT-60.0°C. *II-5*: F-5’-AGCAATGAGACGATGAGGCTT-3’, R-5’-CCCCCACGGACAGTTTGATT-3’; AT-62.4°C. IL-4: F-5’-CCATATCCACGGATGCGACA-3’, R-5’-CTGTGGTGTTCTTCGTTGCTG-3’; AT-60.0°C. *Chil3*: F-5′-GAAGGAGCCACTGAGGTCTG-3′, R-5′-GAGCCACTGAGCCTTCAAC-3′; AT-60.0°C. *Arg1*: F-5′-CAGAAGAATGGAAGAGTCAG-3′, R-5′-CAGATATGCAGGGAGTCACC-3′; AT-60.0°C. *Muc2*: F-5’-GCTGACGAGTGGTTGGTGAATG-3’, R-5’-GATGAGGTGGCAGACAGGAGAC-3’; AT: 60.0°C *Gapdh*: F-5’-AAC TTT GGC ATT GTG GAA GG-3’, R-5’-ACA CAT TGG GGG TAG GAA CA–3’; AT: 62.4°C.

### Preparation of single-cell suspension and flow cytometry

Cecal tissues were cut longitudinally and washed into buffer 1 (HBSS with 25 mM HEPES and 5% FBS). Epithelial cells were separated from the lamina propria by incubating tissue in buffer 2 (HBSS with 15 mM HEPES, 5 mM EDTA, 10% FBS, and 1 mM DTT) for 40 minutes at 37°C with gentle shaking. Then, the lamina propria was dissected into small pieces with scissors and incubated in digestion buffer (RPMI 1640 containing 0.17 mg/mL Liberase TL (Roche, Basel, Switzerland) and 30 µg/mL DNase (Sigma-Aldrich, St. Louis, MO)) for 30 minutes at 37°C with gentle shaking. Single-cell suspensions were made by passing the digested tissue through a 100 µm cell-strainer followed by a 40 µm cell-strainer. Cells were pelleted by centrifuging at 500 x *g* for 5 minutes and reconstituted in FACS buffer (2% FBS in PBS). For flow cytometry, 1×10^6^ cells/sample were incubated with TruStain fcX (BioLegend, San Diego, CA)) for 5 minutes at room temperature, followed by incubation with LIVE/DEAD Fixable Aqua (Life Technologies, Carlsbad, CA) for 30 minutes at 4°C. Cells were washed with FACS buffer twice before staining with monoclonal antibodies targeted to cell surface markers, followed by incubation for 45 minutes at 4°C. Immune cells were detected by flow cytometry using an LSR Fortessa cytometer (BD Biosciences, San Jose, CA). For transcription factor staining, cells were fixed and permeabilized using Foxp3/transcription factor staining buffer (eBioscience, San Diego, CA) before staining with antibodies. Data were analyzed via FlowJo software. The following monoclonal antibodies were used for staining: CD206 (141716), F4/80 (123133), CD19 (115533), CD5 (100623), CD3 (100327), FcεRIα (134317), CD11c (117327), CD90 (105305), CD11b (101215) Ly6C (128005), CD45 (103115), Ly6G (127617), Tbet (505839), from Biolegend (San Diego, CA); ST2 (12-9333-82), RoryT (17-6981-82) from eBiosciences (San Diego, CA); GATA3 (L50-823), SiglecF (552126), CD127 (566377) from BD Bioscience (San Jose, CA).

### ILC2 isolation, expansion, and adoptive transfer

WT C57BL/6J mice were administered 0.75 µg IL-33 for 4 days. Mouse colons, spleens, and mesenteric lymph nodes were collected, and single-cell suspensions were made. ILCs were enriched using mouse Lineage Cell Depletion Kit (Milteny Biotec, Bergisch Gladbach, Germany). Enriched cells were *in vitro* expanded towards ILC2s for 4 days by treating with IL-2 (10 ng/ml), IL-7 (10 ng/ml), and IL-33 (50 ng/ml) ^15^. Lin-CD45+CD127+CD90+CD25+ST2+ ILC2s were then collected by flow-sorting followed by intraperitoneal injection of 5×10^5^ cells or vehicle per mouse.

### Transcriptome microarray

Colon biopsies were collected from eight adult patients who came to the International Centre for Diarrhoeal Diseases Research, Bangladesh with *E. histolytica* infection. Control samples were collected from the same individuals after 60 days of primary infections during recovery. Microarray data were stored in NCBI’s Gene Expression Omnibus (GEO) (http://www.ncbi.nlm.nih.gov/geo/) ^18^. The volcano plot was generated using the DESeq2 package on R ^45^. Consensus PathDB database was used to perform pathway enrichment analysis ^46^. Microarray data on mouse colonic tissue are accessible through GEO accession number GSE43372 ^47^.

### Eosinophil depletion

For eosinophil depletion, IL-33 treated mice received 40 µg anti-SiglecF (clone 238047) or IgG2a isotype control (clone 54447). Successful depletion of eosinophils was confirmed by flow cytometry.

### Statistical analysis

Fisher’s exact test was used to compare the infection rate between groups. Student’s t-test or the Mann-Whitney U nonparametric test were used to make comparisons between two groups. ANOVA was used to compare multiple groups. All the tests were done using GraphPad Prism software.

## Supporting information

Suppelementary Figure

## Acknowledgements

The authors would like to thank the University of Virginia Flow cytometry, Histology, and Biorepository and Tissue Research Facility cores for technical support. We want to thank Drs. Thomas J. Braciale, Janet V. Cross, James E. Casanova, and Chance John Luckey for helpful discussions. The authors would also like to thank the Mann, Ramakrishnan, Marie, and Moonah laboratories for helpful discussions. This work was funded by the National Institutes of Health (NIH) grant R37AI026649 to W.A.P. M.J.U. was also supported by the UVA Robert Wagner Fellowship.

## Author contributions

M.J.U. and W.A.P. designed experiments. M.J.U., J.L., S.L.B., N.O., B.T., and M.A. performed all the experiments. J.L. and S.L.B. gave important advice about designing the experiments. M.J.U. wrote the original draft and W.A.P., J.L., and B.T. reviewed and edited the manuscript. P.K. contributed to analyzing transcriptomic data. S.L.B., A.L.F., and A.N.D., helped to design flow cytometry panels, and A.L.F provided invaluable advice in processing colonic tissue to isolate single-cell suspensions. All the authors reviewed the manuscript.

## Figure Legends

**Supplementary figure 1:**
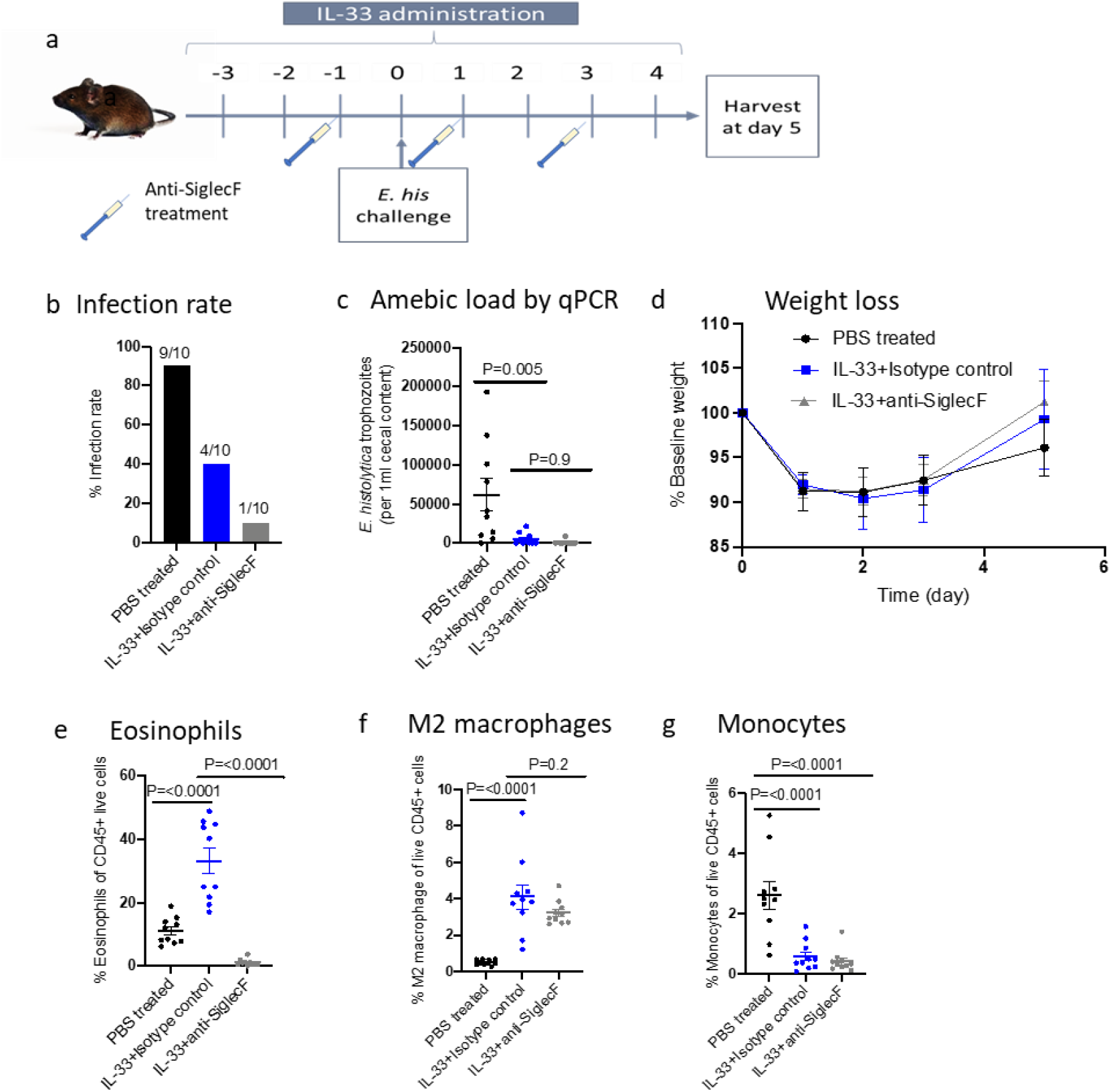
IL-33 mediated protection from amebic colitis is not mediated by eosinophils. **a-g** CBA/J mice were treated with 0.75 µg of IL-33 or PBS for 8 days. IL-33 treated groups were administered 3 doses of anti-SiglecF antibody or isotype control. **a** Experimental outline. **b** Infection rate by amebic culture. **c** Amebic load by qPCR. **d** Weight loss. **e-f** Percentage of eosinophils, CD206+ macrophages, and monocytes in cecal tissue. **b-g** n=10, per group. Statistical significance was determined by one-way ANOVA. Error bars indicate SEM.

## References

1. Liu, L. et al. Global, regional, and national causes of child mortality: an updated systematic analysis for 2010 with time trends since 2000. Lancet 379, 2151–2161 (2012).

2. Mondal, D., Petri, W. A., Sack, R. B., Kirkpatrick, B. D. & Haque, R. Entamoeba histolytica-associated diarrheal illness is negatively associated with the growth of preschool children: evidence from a prospective study. Trans. R. Soc. Trop. Med. Hyg. 100, 1032–1038 (2006).

3. Lozano, R. et al. Global and regional mortality from 235 causes of death for 20 age groups in 1990 and 2010: a systematic analysis for the Global Burden of Disease Study 2010. Lancet 380, 2095– 2128 (2012).

4. Irusen, E. M., Jackson, T. F. H. G. & Simjee, A. E. Asymptomatic Intestinal Colonization by Pathogenic Entamoeba histolytica in Amebic Liver Abscess: Prevalence, Response to Therapy, and Pathogenic Potential. Clin. Infect. Dis. 14, 889–893 (1992).

5. Uddin, M. J., Leslie, J. L. & Petri, W. A. Host Protective Mechanisms to Intestinal Amebiasis. Trends Parasitol. 37, 165–175 (2021).

6. Ralston, K. S. et al. Trogocytosis by Entamoeba histolytica contributes to cell killing and tissue invasion. Nature 508, 526–530 (2014).

7. Bansal, D. et al. An ex-vivo Human Intestinal Model to Study Entamoeba histolytica Pathogenesis. PLoS Negl. Trop. Dis. 3, e551 (2009).

8. Watanabe, K. et al. Microbiome-mediated neutrophil recruitment via CXCR2 and protection from amebic colitis. PLOS Pathog. 13, e1006513 (2017).

9. Ghadiriani, E. & Bout, D. T. In vitro Killing of Entamoeba histolytica trophozoites by interferon-γ-activated mouse macrophages. Immunobiology 176, 341–353 (1988).

10. Petri, W. A. et al. CORRELATION OF INTERFERON-γ PRODUCTION BY PERIPHERAL BLOOD MONONUCLEAR CELLS WITH CHILDHOOD MALNUTRITION AND SUSCEPTIBILITY TO AMEBIASIS. Am. J. Trop. Med. Hyg. 76, 340–344 (2007).

11. Mondal, D. et al. Association between TNF-α and Entamoeba histolytica Diarrhea. Am. J. Trop. Med. Hyg. 82, 620–625 (2010).

12. Noor, Z. et al. Role of Eosinophils and Tumor Necrosis Factor Alpha in Interleukin-25-Mediated Protection from Amebic Colitis. MBio 8, (2017).

13. Moltke, J. von, Ji, M., Liang, H.-E. & Locksley, R. M. Tuft-cell-derived IL-25 regulates an intestinal ILC2–epithelial response circuit. Nature 529, 221–225 (2016).

14. Tu, L. et al. IL-33-induced alternatively activated macrophage attenuates the development of TNBS-induced colitis. Oncotarget 8, 27704–27714 (2017).

15. Frisbee, A. L. et al. IL-33 drives group 2 innate lymphoid cell-mediated protection during Clostridium difficile infection. Nat. Commun. 10, 2712 (2019).

16. Monticelli, L. A. et al. IL-33 promotes an innate immune pathway of intestinal tissue protection dependent on amphiregulin–EGFR interactions. Proc. Natl. Acad. Sci. 112, 10762–10767 (2015).

17. Barlow, J. L. et al. IL-33 is more potent than IL-25 in provoking IL-13–producing nuocytes (type 2 innate lymphoid cells) and airway contraction. J. Allergy Clin. Immunol. 132, 933–941 (2013).

18. Peterson, K. M. et al. The expression of REG 1A and REG 1B is increased during acute amebic colitis. Parasitol. Int. 60, 296–300 (2011).

19. Houpt, E. R. et al. The Mouse Model of Amebic Colitis Reveals Mouse Strain Susceptibility to Infection and Exacerbation of Disease by CD4 + T Cells. J. Immunol. 169, 4496–4503 (2002).

20. Schmitz, J. et al. IL-33, an Interleukin-1-like Cytokine that Signals via the IL-1 Receptor-Related Protein ST2 and Induces T Helper Type 2-Associated Cytokines. Immunity 23, 479–490 (2005).

21. Hayakawa, H., Hayakawa, M., Kume, A. & Tominaga, S. Soluble ST2 Blocks Interleukin-33 Signaling in Allergic Airway Inflammation. J. Biol. Chem. 282, 26369–26380 (2007).

22. Humphreys, N. E., Xu, D., Hepworth, M. R., Liew, F. Y. & Grencis, R. K. IL-33, a Potent Inducer of Adaptive Immunity to Intestinal Nematodes. J. Immunol. 180, 2443–2449 (2008).

23. Yamaguchi, Y. et al. Purified interleukin 5 supports the terminal differentiation and proliferation of murine eosinophilic precursors. J. Exp. Med. 167, 43–56 (1988).

24. Waddell, A., Vallance, J. E., Hummel, A., Alenghat, T. & Rosen, M. J. IL-33 Induces Murine Intestinal Goblet Cell Differentiation Indirectly via Innate Lymphoid Cell IL-13 Secretion. J. Immunol. 202, 598–607 (2019).

25. Lidell, M. E., Moncada, D. M., Chadee, K. & Hansson, G. C. Entamoeba histolytica cysteine proteases cleave the MUC2 mucin in its C-terminal domain and dissolve the protective colonic mucus gel. Proc. Natl. Acad. Sci. 103, 9298–9303 (2006).

26. Chadee, K., Petri, W. A., Innes, D. J. & Ravdin, J. I. Rat and human colonic mucins bind to and inhibit adherence lectin of Entamoeba histolytica. J. Clin. Invest. 80, 1245–1254 (1987).

27. Schiering, C. et al. The alarmin IL-33 promotes regulatory T-cell function in the intestine. Nature 513, 564–568 (2014).

28. Malik, A. et al. IL-33 regulates the IgA-microbiota axis to restrain IL-1α–dependent colitis and tumorigenesis. J. Clin. Invest. 126, 4469–4481 (2016).

29. Kobori, A. et al. Interleukin-33 expression is specifically enhanced in inflamed mucosa of ulcerative colitis. J. Gastroenterol. 45, 999–1007 (2010).

30. Liew, F. Y., Girard, J.-P. & Turnquist, H. R. Interleukin-33 in health and disease. Nat. Rev. Immunol. 16, 676–689 (2016).

31. Huang, J. et al. Hyperactivity of Innate Immunity Triggers Pain via TLR2-IL-33-Mediated Neuroimmune Crosstalk. Cell Rep. 33, 108233 (2020).

32. Wong-Baeza, I. et al. The Role of Lipopeptidophosphoglycan in the Immune Response to Entamoeba histolytica. J. Biomed. Biotechnol. 2010, 1–12 (2010).

33. Stojkovic, S. et al. Tissue factor is induced by interleukin-33 in human endothelial cells: a new link between coagulation and inflammation. Sci. Rep. 6, 25171 (2016).

34. Ishihara, K. & Hirano, T. IL-6 in autoimmune disease and chronic inflammatory proliferative disease. Cytokine Growth Factor Rev. 13, 357–368 (2002).

35. Klion, A. D. & Nutman, T. B. The role of eosinophils in host defense against helminth parasites. J. Allergy Clin. Immunol. 113, 30–37 (2004).

36. Eberl, G., Colonna, M., Santo, J. P. D. & McKenzie, A. N. J. Innate lymphoid cells: A new paradigm in immunology. Science (80-.). 348, aaa6566–aaa6566 (2015).

37. Moro, K. et al. Innate production of TH2 cytokines by adipose tissue-associated c-Kit+Sca-1+ lymphoid cells. Nature 463, 540–544 (2010).

38. Nakamura, R. et al. Group 2 Innate Lymphoid Cells Exacerbate Amebic Liver Abscess in Mice. iScience 23, 101544 (2020).

39. Schuijs, M. J. et al. ILC2-driven innate immune checkpoint mechanism antagonizes NK cell antimetastatic function in the lung. Nat. Immunol. 21, 998–1009 (2020).

40. Gieseck, R. L., Wilson, M. S. & Wynn, T. A. Type 2 immunity in tissue repair and fibrosis. Nat. Rev. Immunol. 18, 62–76 (2018).

41. Mchedlidze, T. et al. Interleukin-33-Dependent Innate Lymphoid Cells Mediate Hepatic Fibrosis. Immunity 39, 357–371 (2013).

42. Zeis, P. et al. In Situ Maturation and Tissue Adaptation of Type 2 Innate Lymphoid Cell Progenitors. Immunity 53, 775–792.e9 (2020).

43. Diamond, L. S., Harlow, D. R. & Cunnick, C. C. A new medium for the axenic cultivation of Entamoeba histolytica and other Entamoeba. Trans. R. Soc. Trop. Med. Hyg. 72, 431–432 (1978).

44. Buonomo, E. L. et al. Microbiota-Regulated IL-25 Increases Eosinophil Number to Provide Protection during Clostridium difficile Infection. Cell Rep. 16, 432–443 (2016).

45. Love, M. I., Huber, W. & Anders, S. Moderated estimation of fold change and dispersion for RNA-seq data with DESeq2. Genome Biol. 15, 550 (2014).

46. Kamburov, A., Stelzl, U., Lehrach, H. & Herwig, R. The ConsensusPathDB interaction database: 2013 update. Nucleic Acids Res. 41, D793–D800 (2013).

47. Mackey-Lawrence, N. M. et al. Effect of the Leptin Receptor Q223R Polymorphism on the Host Transcriptome following Infection with Entamoeba histolytica. Infect. Immun. 81, 1460–1470 (2013).

