## Supplementary material for "The IL-33-ILC2 pathway protects from amebic colitis": Suppelementary Figure

**Supplementary figure 1: IL-33 mediated protection from amebic colitis is not mediated by eosinophils.**

**a-g** CBA/J mice were treated with 0.75 µg of IL-33 or PBS for 8 days. IL-33 treated groups were administered 3 doses of anti-SiglecF antibody or isotype control. **a** Experimental outline. **b** Infection rate by amebic culture. **c** Amebic load by qPCR**. d** Weight loss. **e-f** Percentage of eosinophils, CD206+ macrophages, and monocytes in cecal tissue. **b-g** n=10, per group. Statistical significance was determined by one-way ANOVA. Error bars indicate SEM.

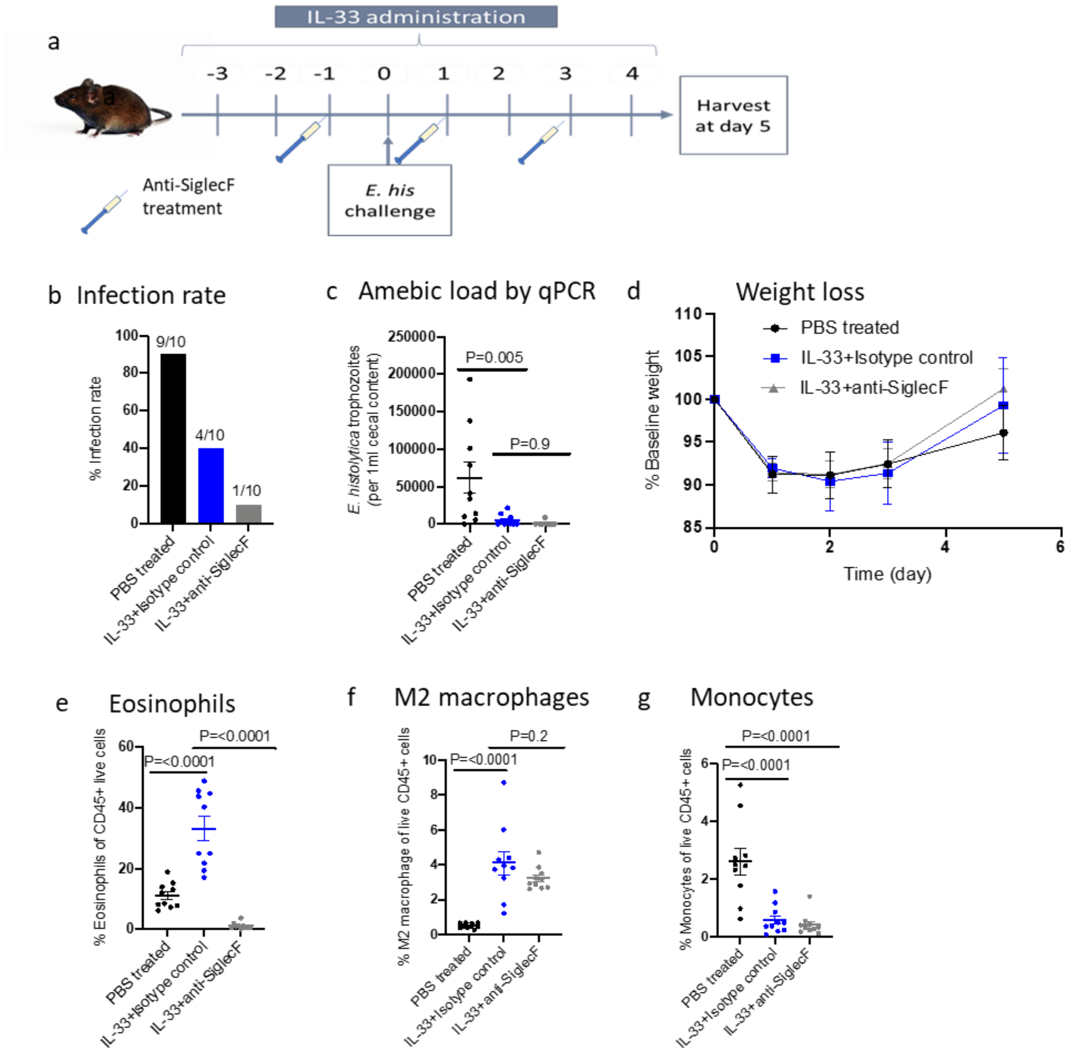
